# Recurrent LINE 1 exonization drives transcriptome remodelling in NSCLC

**DOI:** 10.64898/2026.04.22.720055

**Authors:** Ankita Subhadarsani Parida, Arun Kumar, Bhavana Tiwari

**Affiliations:** Department of Biological Sciences, Indian Institute of Science Education and Research (IISER) Berhampur, Odisha, India

**Keywords:** LINE-1/L1, NSCLC, RNA-seq, Exonixation

## Abstract

The only autonomously active transposable elements in the human genome are Long interspersed nuclear element-1 (LINE-1) elements. These elements are known to play an important role in changing the transcriptome. LINE-1 sequences affect gene regulation during post-transcription processing, along with their established role in retrotransposition. Exonization is one mechanism where the LINE-1 integrated genome undergoes alternative splicing to produce new isoforms of transcripts.

Our work mainly highlights the effect of LINE-1 associated exonization, focusing on the formation of isoforms of transcripts. Using Non-small cell lung cancer (NSCLC) as a model, we conducted a detailed transcriptome study that combines splice junction profiling with gene expression data. Our results show that LINE-1 sequences are often included as exons in host transcripts, leading to the formation of new exons and their various isoforms. The events are validated by solid splice junction evidence that proves the reliability and reproducibility. In particular, it was observed that repetitive analyses revealed certain LINE-1 exonization events that were consistent. The finding indicates that LINE-1 act as recurrent sources of splice ready sequences. Though exonizations do not necessarily affect the total expression levels of genes, our study reveals that they certainly contribute to transcript diversity. The diversity of isoforms generated potentially contributes to the effects of gene function. This study is limited to NSCLC, but it is likely that the exonizations events play a crucial role in the altering RNA diversity in cancers. Therefore the study elucidates new insights into how transposable elements modify gene structure and function during cancer development.

## Introduction

The only autonomously active family of non-long terminal repeat (non-LTR) transposable elements in the human genome is long interspersed nuclear element-1 (LINE-1/L1), which makes up around 17% of the human genome and has ∼500,000 copies across it (Ramos et al., 2021). A portion of these elements are still capable of retrotransposition through an RNA intermediate and are transcriptionally competent (Elbarbary et al., 2016; Gebrie, 2023). LINE-1 elements have a variety of regulatory impacts on gene expression outside of their typical involvement in insertional mutagenesis, especially at the level of transcription and RNA processing (Beck et al., 2011; Elbarbary et al., 2016; Okorokova, 2025). Exonization is a process, whereby genes acquire new exons from non-protein-coding (Schmitz & Brosius, 2011). In LINE-1 sequences these intronic regions are co-opted into mature transcripts through alternative splicing to produce LINE-1 derived exons or chimeric transcripts, is a well-established fact by which LINE-1 affects gene regulation (Gebrie, 2023; Khalid et al., 2018; *Line1*, n.d.; Mir, 2015). LINE-1 antisense promoter has the ability of to push transcription into nearby genic areas by promoting transcript diversification (Han & Boeke, 2005; Xu et al., 2023). Since aberrant LINE-1 activation has been connected to changed gene function and carcinogenic phenotypes in cancer, these mechanisms are becoming more widely studied (Lavia et al., 2022; McKerrow et al., 2022; Pessoa et al., 2026; Xiao-Jie et al., 2016). In healthy somatic tissues the mechanisms of LINE-1 activity is strictly controlled by promoter methylation, transcriptional repression, and premature polyadenylation (Thayer et al., 1993; Takai et al., 2000). However, these limitations are often deregulated in cancer, leading to increased transcriptional output and LINE-1 reactivation. Although LINE-1 retrotransposition has been thoroughly studied at the DNA level, but its effects at the RNA level, especially exonization and its role in transcriptome remodelling is under explored (Kemp & Longworth, 2015; Mendez-Dorantes & Burns, 2023; Sciamanna et al., 2016). NSCLC, which accounts for the majority of Lung cancer cases and remains a leading cause of Worldwide death (Barta et al., n.d.; Gridelli et al., 2015; Hendriks et al., 2024; Leiter et al., 2023; Molina et al., 2008; Sharma, 2022). In addition to well characterized genetic alteration, this exhibits broad dysregulation of RNA processing, including alternative splicing (Bradley & Anczuków, 2023; Yan et al., 2023). Emerging evidence suggests that transposable elements contribute to transcriptomic diversity, but the role of LINE-1 mediated exonization in cancer remains underexplored (Burns, 2017; Ponomaryova et al., 2020),Systematic identification of such events is technically challenging due to the repetitive nature of LINE-1 sequences and the limitations of short read RNA sequencing in accurately resolving splice junctions (Baumann et al., 2025). Consequently, prior studies have largely focused on isolated exonization events or DNA-level insertions, and comprehensive transcriptome-wide analyses across patient cohorts remain limited. not received enough attention despite notable progress in finding oncogenic drivers. Because LINE-1 sequences are repetitive and short-read RNA sequencing cannot reliably resolve splice junctions, systematic detection of LINE-1 exonization events is technically difficult. As a result, previous research has mostly concentrated on single events or DNA level insertions, and thorough transcriptome-wide assessments across patient cohorts are still scarce.

Here, we performed a transcriptome wide investigation to systematically characterize LINE-1 mediated exonization in NSCLC. Using matched tumor and normal samples (Seo et al., 2012), we integrated differential gene expression analysis, splice junction profiling, and recurrence mapping across patient samples. Our analysis identified widespread incorporation of LINE-1 derived sequences into host transcripts, supported by high confidence splice junction evidence, indicating bona fide alternative splicing events. We studied the link between host gene expression and LINE-1 activity. A significant correlation was observed in the case of the presence of LINE-1 with genes exhibiting exonizations. Chromosome mapping for exonization events revealed variance among the samples. This illustrates that LINE-1 mediated transcript remodeling can vary from patient to patient. Several novel cryptic exonization events were observed in genes that showed inconsistent expression. In order to validate our results and highlight the differences in the splicing patterns of matched normal and tumor samples, we used the integrative genomics viewer (IGV). Moreover, we observed several canonical donor and acceptor sites associated with LINE-1-derived exons by splice junction architecture analysis. The recurrence of exonization events in several patients implies that LINE-1 elements could act as a potential source of splicing ready sequence in cancers. The importance of this study is further highlighted considering the fact that the study is only focused on NSCLC. LINE-1 elements play an important role in alternative splicing. They help in generating isoforms of genes that increase the transcript diversity without affecting the overall gene expression level. Thus, the above discussion reveals the significance of LINE-1 exonization in the alteration of cancer genes’ transcripts.

## Results

### Transcriptomic dysregulation and association with LINE-1 expression in NSCLC

We performed differential gene expression analysis on paired tumor (n = 12) and normal tissue samples (n = 12) to fully describe the transcriptional landscape of NSCLC and assess its relationship to LINE-1 activity. Our work revealed substantial reprogramming of the transcriptome in tumor tissues. 5,492 genes were significantly increased in tumors and 1,668 genes were downregulated in comparison to matched normal samples (**Fig. 1A**). The distribution of differentially expressed genes in NSCLC displayed a substantial bias towards upregulation rather than balanced bidirectional changes, indicating a global tendency of transcriptional activation. Unsupervised hierarchical clustering of these genes produced a heatmap that demonstrated a clear and robust segregation between tumor and matched normal samples (**Fig. 1B**). The high expression of the transcriptional differences found is highlighted by this special partitioning, which shows that the significant changes in gene expression are repeated throughout the cohort and cannot be explained by technical noise or random fluctuation. The transcriptional changes in the tumor samples that differ from the molecular indicators of healthy lung tissue is highlighted by the clustering pattern. Significantly, thousands of genes rather than a small number contributed to the separation, indicating that NSCLC is characterized by a widespread and well-coordinated reconfiguration of gene expression networks. This consistency across all samples offers strong proof that NSCLC biology is characterized by transcriptional dysregulation. Next, the expression profiles of key signaling molecules and oncogenes associated with NSCLC were examined. Several well-known oncogenic drivers, such as BRAF, CTNNB1, EGFR, KRAS, MET, NRAS, and PIK3CA, were found to have higher expression in tumor samples compared to matched normal tissues, according to a comparative analysis employing boxplot visualization (**Fig. 1C**). Furthermore, elevated expression was noted for MIGA1, LSR, TADA1, and WASHC3, which have developing or context-dependent functions in cancer biology despite not being traditional oncogenic drivers.

**Figure 1.**
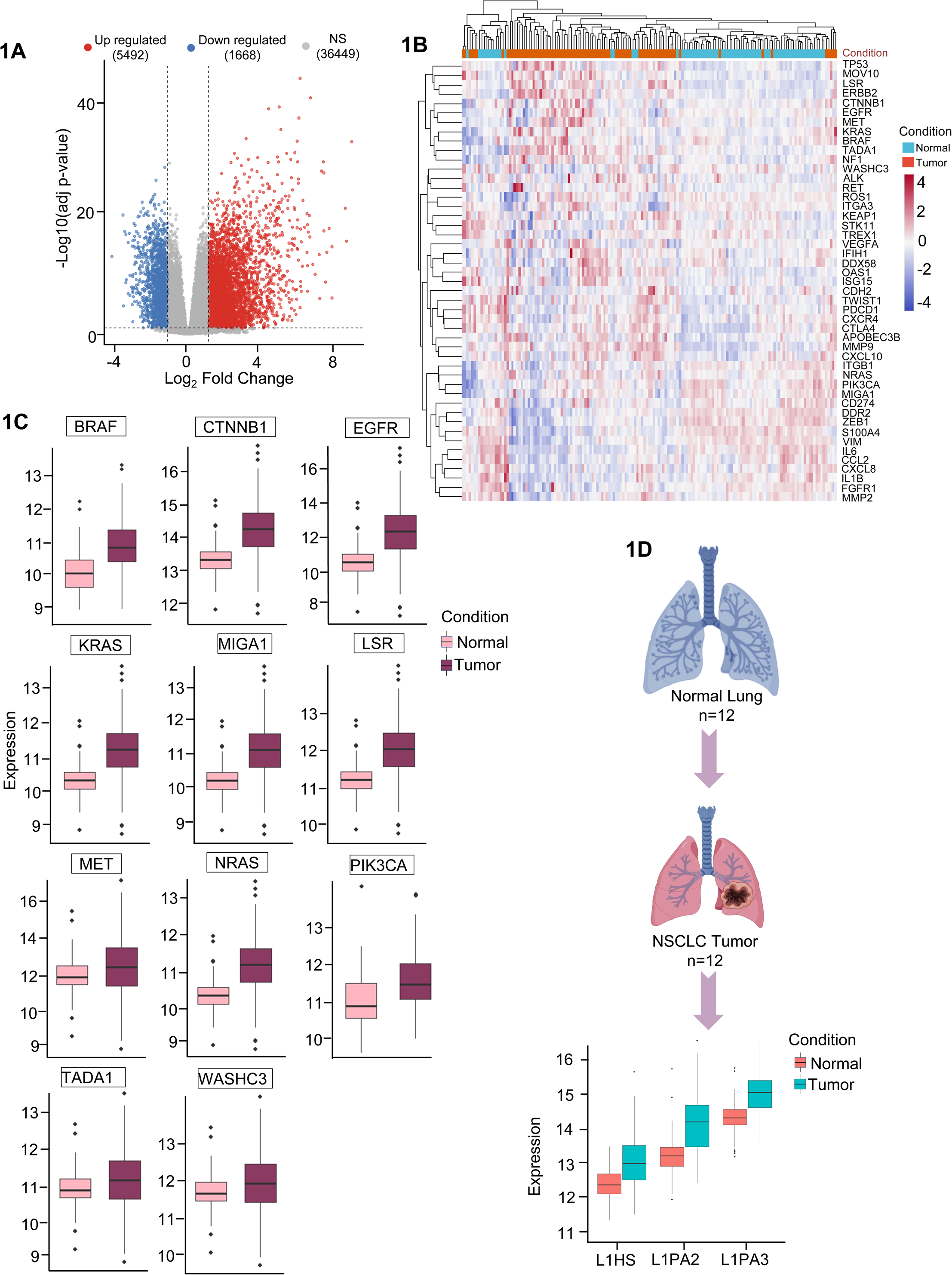
Transcriptomic dysregulation and association with LINE-1 expression in NSCLC. **(A)** Volcano plot of differential gene expression between tumor and normal samples generated using DESeq2. The x-axis represents log₂ fold change and the y-axis shows - log₁₀ adjusted p-values (Benjamini-Hochberg correction). Genes were considered significantly differentially expressed at |log₂FC| ≥ 1 and adjusted p-value (FDR) < 0.05. Upregulated genes are shown in red, downregulated genes in blue, and non-significant genes in gray. Dashed lines indicate the applied thresholds. **(B)** Heatmap of significantly differentially expressed genes identified by DESeq2 (FDR < 0.05), clustered using hierarchical clustering (Euclidean distance, complete linkage). Rows represent genes and columns represent samples. Expression values are z-score normalized across samples. **(C)** Boxplots showing the expression of established oncogenes (BRAF, CTNNB1, EGFR, KRAS, MET, NRAS, PIK3CA) together with emerging or less-characterized cancer relevant genes (MIGA1, LSR, TADA1, WASCH3) in normal and tumor samples. Statistical significance between groups was assessed using a two-sided Wilcoxon rank-sum test. Center lines indicate medians, boxes represent interquartile ranges (IQR), and whiskers denote 1.5× IQR. **(D)** Graphical summaries of sample distributions for normal and tumor groups are shown alongside boxplots comparing the expression of LINE-1 subfamilies (L1HS, L1PA2, L1PA3). Statistical significance was determined using a two-sided Wilcoxon rank-sum test, with increased expression observed in tumor samples.

All of these findings point to a widespread activation of oncogenic signalling pathways in NSCLC, along with the overexpression of other genes that might be involved in transcriptome remodelling and cellular processes linked to tumors. These results are consistent with the activation of traditional carcinogenic pathways and confirm the biological importance of the dataset. The rise in these drivers further validates the dataset as typical of NSCLC biology and highlights the convergence of transcriptional activity with known carcinogenic pathways. To better characterize transposable element activity, we assessed the expression of LINE-1 families in many samples. Tumor samples exhibited considerably higher expression of several LINE-1 subfamilies, such as L1HS, L1PA2, and L1PA3 elements, compared to normal tissue (**Fig. 1D**). This increase was statistically significant and consistent across the sample, suggesting that NSCLC transcriptomes frequently exhibit LINE-1 element activation. A basis for investigating the molecular connections between transposable element activity and carcinogenic signaling is provided by the coordinated overexpression of LINE-1 families, which emphasizes their possible significance as drivers or modulators of transcriptional dysregulation in NSCLC.

### LINE-1 associated exonization and its relationship with oncogenic gene expression in NSCLC

We conducted gene-wise correlation analysis between L1 family expression and certain oncogenes and cancer-associated genes in NSCLC samples in order to better explore the relationship between LINE-1 activity and host gene expression. A number of important oncogenic drivers, such as BRAF, EGFR, KRAS, MET, PIK3CA, CTNNB1, NRAS, MIGA1, and TADA1 showed moderate to high positive correlations with LINE1 expression, suggesting a possible connection between oncogenic transcriptional programs and LINE-1 activation (**Fig.2A**). Additionally, LSR, TADA1 and WASHC3 showed moderate positive associations, suggesting that LINE-1 activity may also influence non classical cancer associated genes. Genes like, on the other hand, showed minimal associations, suggesting that the effects of LINE-1 expression are not consistent across all genes. Notably, a subset of genes demonstrated inverse correlations like EMT, highlighting potential repressive or compensatory regulatory effects. Significantly, these findings imply that LINE-1-associated gene regulation is locus-specific and context-dependent, where the result transcriptional activation or repression likely depends on the genomic location of LINE-1 elements and how they interact with neighboring genes (e.g, exonization, promoter activity, or splice site contribution). A hypothesis in which LINE-1 contributes to selective and context-dependent transcriptome remodelling in NSCLC is supported by these data, which collectively show a gene-specific and diverse link between LINE-1 activity and host gene expression. We examined the positional effects of intronic LINE-1 sequence on splicing patterns in order to better understand the molecular mechanism of LINE-1-mediated transcript remodelling. Canonical exonisation as well as the different isotype of LINE1 dependent exonisation in which LINE-1 elements found within intronic regions can act as sources of cryptic splice sites, resulting in exonization and the creation of novel transcript isoforms as shown in the Schematic representation (**Fig. 2B**). Significantly, these findings imply that LINE-1 associated gene regulation is locus-specific and context-dependent, where the result transcriptional activation or repression likely depends on the genomic location of LINE-1 elements and how they interact with neighboring genes. A hypothesis in which LINE-1 contributes to selective and context dependent transcriptome remodelling in NSCLC is supported by these data, which collectively show a gene-specific and diverse link between LINE-1 activity and host gene expression. We examined the positional effects of intronic LINE-1 insertions on splicing patterns in order to better understand the molecular mechanism of LINE-1 mediated transcript remodelling. Moreover, isoform diversification can be aided by the incorporation of LINE-1 derived exonic sequences, which can modify transcript architecture without necessarily upsetting the entire gene structure. Such data support a hypothesis suggesting that LINE-1 sequences act as a depot of potential splice sites that could be used in suitable conditions, particularly in the case of cancer when there are frequent disruptions in splicing regulation. Quantitative analysis within the whole group demonstrated a significant excess of LINE-1 exonizations in tumor samples as compared with normal lung tissue (median fold-change, two-sided Wilcoxon rank-sum test, P < 0.001) (**Fig. 2C**). The increased content of such events indicates activation of cryptic splice sites in cancer. We estimated the number of exonization events per patient within the NSCLC group to analyze the distribution of LINE-1 mediated exonizations among samples (**Fig. 2D**). There was considerable variation in exonization frequencies between different samples, indicating that some samples, such as ERR164502 and ERR164495, experienced a greater level of LINE-1 exonization than other samples. The variation indicates that different cancers contain different amounts of LINE-1 exonizations. Nevertheless, LINE-1 exonization occurred in all tissues examined, demonstrating the importance of LINE-1 function in NSCLC.

**Figure 2.**
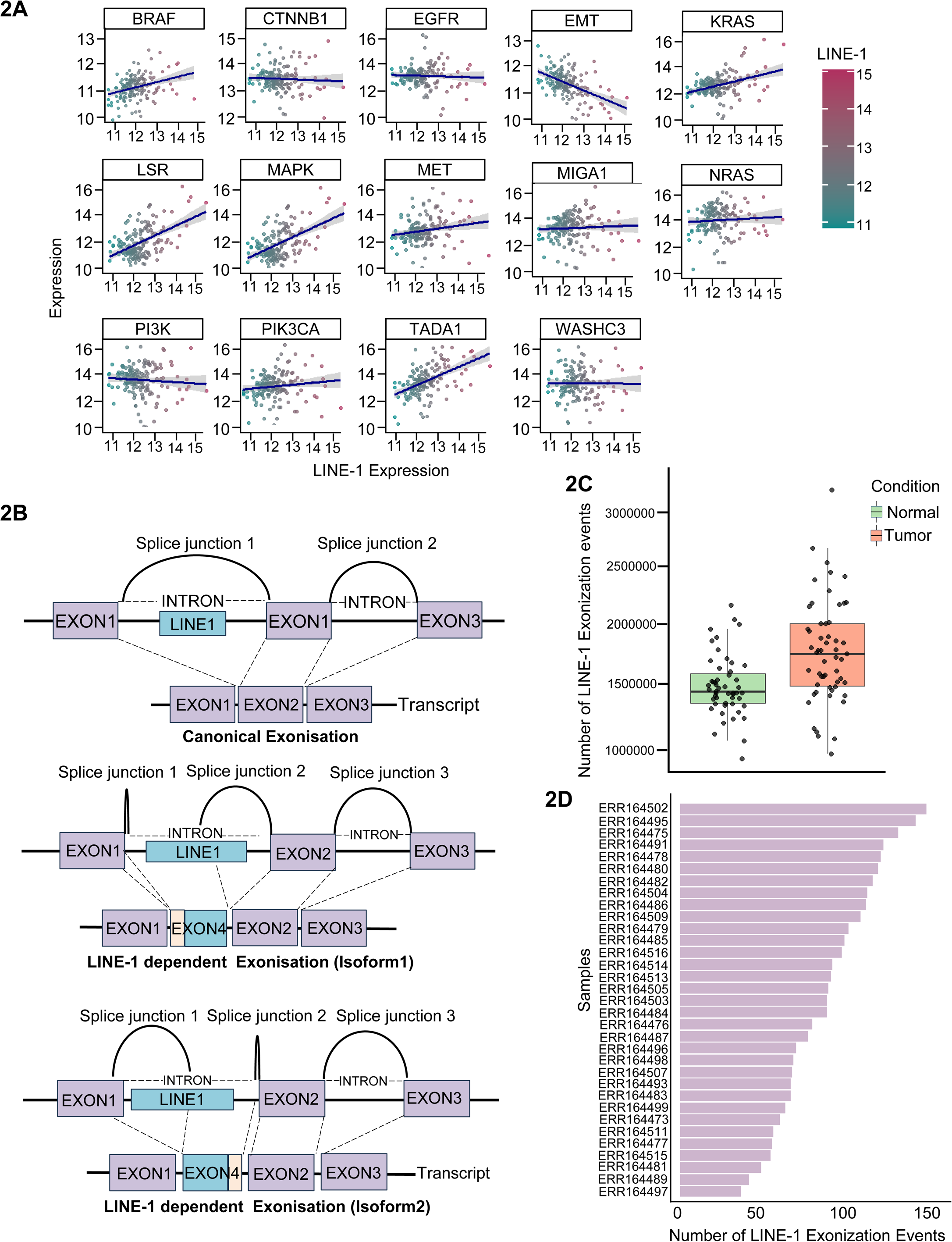
LINE-1 associated exonization and its relationship with oncogenic gene expression in NSCLC. **(A)** Scatter plots showing the relationship between LINE-1 expression and expression of cancer-associated genes (BRAF, CTNNB1, EGFR, EMT markers, KRAS, LSR, MAPK, MET, MIGA1, NRAS, PI3K, PIK3CA, TADA1, and WASHC3). Each point represents an individual sample, colored by LINE-1 expression level. Linear regression lines with 95% confidence intervals are shown. Correlations were assessed using Pearson correlation analysis, and trends indicate both positive and negative associations between LINE-1 activity and oncogenic pathways. **(B)** Schematic representation of LINE-1 mediated exonization mechanisms. Top: Canonical splicing showing standard exon joining (Exon1-Exon2-Exon3) with intron removal. Middle and bottom: LINE-1 dependent exonization events, where intronic LINE-1 sequences introduce novel splice donor and/or acceptor sites, resulting in inclusion of cryptic exons (e.g., Exon4) into the mature transcript. Multiple isoforms can arise depending on splice junction usage. **(C)** Boxplot comparing the number of LINE-1 exonization events between normal and tumor samples. Tumor samples exhibit a higher number of exonization events relative to normal tissues. Statistical significance was assessed using a two-sided Wilcoxon rank-sum test. Center lines represent medians, boxes indicate interquartile range (IQR), and whiskers denote 1.5× IQR. **(D)** Bar plot showing the number of LINE-1 exonization events per sample across the cohort. Each bar represents an individual sample (ERR IDs), highlighting inter-sample variability in LINE-1 mediated exonization and indicating that certain tumor samples exhibit markedly elevated exonization activity.

The idea that LINE-1 mediated exonization is a frequent feature of the NSCLC transcriptome is further supported by the fact that most samples showed a moderate to high number of exonization events, which is significant in all the cancer sample. The observed inter-sample heterogeneity could be due to variations in the splicing machinery among tumors, variations in LINE-1 activation state, or variations in epigenetic regulation. When taken as a whole, these results demonstrate how LINE-1-driven transcriptome remodelling is both patient-specific and widespread, supporting its importance as a major factor in transcriptional complexity in NSCLC.

### Genomic distribution, strand bias, and splice site architecture of LINE-1 exonization events

We created a binary presence absence matrix of exonization events across all samples and genomic locations in order to examine the distribution and recurrence of LINE-1 mediated exonization events throughout the NSCLC cohort. This matrix’s unsupervised hierarchical clustering showed that LINE-1 exonization events were widely detected throughout the population (**Fig. 3A**). The heatmap shows that a number of exonization events are frequently seen in various samples, suggesting the existence of conserved, splice-competent LINE-1 sites. Furthermore, clustering analysis suggests common underlying mechanisms of LINE-1 linked splicing regulation by grouping samples according to shared exonization patterns. Interestingly, LINE-1 exonization events are consistently seen in most samples, which lends credence to the notion that these events are a prevalent and functionally significant aspect of the transcriptome of NSCLC. Further evidence that some LINE-1 sites may preferentially contribute to transcript synthesis comes from the recurrence of particular exonization events. All of these results point to LINE-1 mediated exonization as a common and recurrent process in NSCLC, which contributes to transcriptome remodeling by repeatedly using splice-competent LINE-1 sequences. We measured exonization events resulting from sense and antisense orientations in both NSCLC and normal samples in order to further examine the strand-specific contribution of LINE-1 mediated exonization (**Fig. 3B**). Tumor samples showed a significant increase in exonization events in both sense and antisense orientations in comparison to normal tissues. Significantly, exonization events from the antisense orientation were more common than those from the sense orientation, which is in line with the antisense strand of LINE-1 elements having splice compatible features. All of these results show that LINE-1 mediated exonization is strongly increased in NSCLC and happens in a strand dependent manner with a predominance of antisense driven events. We examined the nucleotide composition around splice donor and acceptor sites connected to LINE-1 generated exons in order to further describe the sequence properties underpinning LINE-1 driven exonization events. Both donor and acceptor junctions had conserved splice site motifs, according to sequence logo analysis (**Fig.3C**). The canonical GT dinucleotide was clearly enriched at the splice donor sites, which is in line with known 5′ splice site consensus sequences.

**Figure 3.**
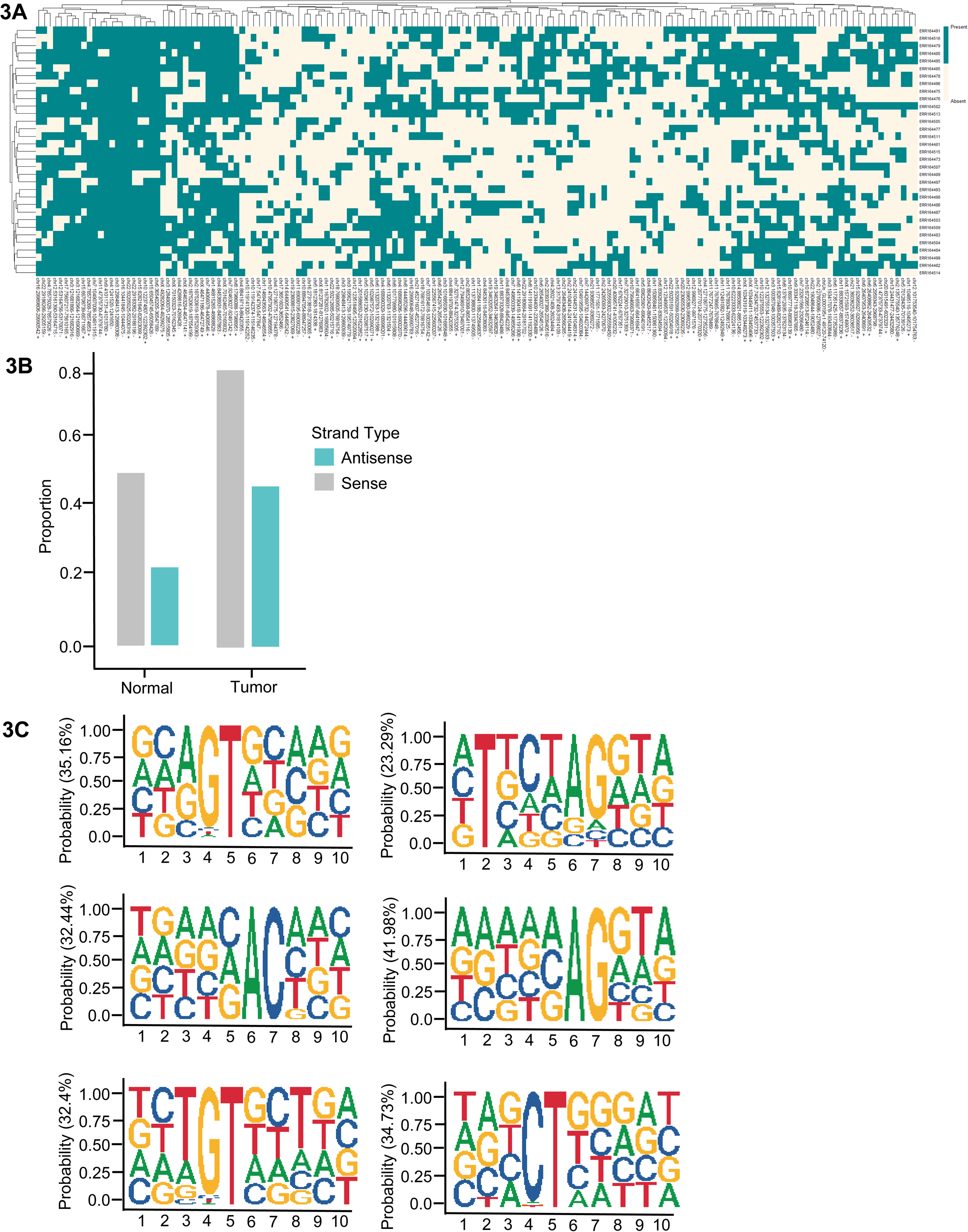
Genomic distribution, strand bias, and splice site architecture of LINE-1 exonization events. (A) Binary heatmap representing the presence or absence of LINE-1 exonization events across samples and loci. Hierarchical clustering highlights recurrent exonization patterns and substantial inter-sample heterogeneity. (B) Strand orientation analysis of exonization events. The proportion of sense and antisense LINE-1 insertions is shown for normal and tumor samples. Tumors exhibit increased exonization across both orientations. (C) Sequence logos of splice sites associated with LINE-1 exonization. Motif analysis of ±5 bp flanking junctions reveals canonical splice signatures, including conserved GT donor and AG acceptor motifs, alongside variable flanking nucleotide preferences. Distinct motif clusters and their relative frequencies (%) indicate multiple sequence contexts capable of supporting LINE-1 driven exonization.

In a similar vein, splice acceptor sites showed a significant upstream polypyrimidine tract and a substantial enrichment of the conventional AG dinucleotide, both of which are signs of functioning 3′ splice site architecture. Significantly, these conserved patterns were found in both sense and antisense LINE-1 derived exonization events, indicating that LINE-1 sequences contain splice compatible regions that the splicing machinery can recognize. Sequence variety was reflected in the little differences in nucleotide content surrounding these core motifs, which preserved crucial splice site characteristics. Together, these findings show that conventional and functional splice signals support LINE-1 mediated exonization, supporting the idea that LINE-1 elements act as a storehouse of cryptic splice sites that can be used to create new exons.

### Genome browser visualization of LINE-1 associated exonization events in NSCLC

We used integrated visualization with IGV to compare splice junctions and read coverage between control and NSCLC samples in order to validate LINE-1 mediated exonization events at the locus level. Clear evidence of cancer-specific splicing changes linked to intronic LINE-1 elements was found by representative analysis at the TADA1, MIGA1, LSR, and WASHC3 locus (**Fig. 4 and 5**). Splice junctions in control samples had consistent junction support and the anticipated transcript structure, adhering to the standard exon-intron design. Tumor samples, on the other hand, showed new splice junctions that overlapped LINE-1 elements along with increased read coverage throughout the region, suggesting that LINE-1 derived sequences were incorporated into mature transcripts. Crucially, the reported exonization event is associated with a LINE-1 locus, where cryptic splice donor and acceptor sites are either absent or used very infrequently in controls but are exploited in cancer cases. leads to the creation of a new exon in the host gene, which is in line with LINE-1 driven exonization. Strong evidence that these events are real alternative splicing events rather than mapping artifacts is also provided by the presence of strong junction reads spanning the LINE-1 region in tumor samples together with altered coverage patterns. Together, these locus-specific validations corroborate our transcriptome-wide results, showing that LINE-1 elements can be exonized into gene architecture and contribute to the diversity of cancer-specific transcript isoforms. We used IGV in conjunction with splice junction mapping and sequence characterisation to conduct locus-specific analysis in order to further clarify at the gene level. Clear evidence of LINE-1 driven exonization events in tumor tissues was found by representative analysis at the WASHC3 gene (**Fig. 5**). RNA sequencing coverage and splice junction visualization revealed the existence of cancer-specific splice junctions that overlapped intronic LINE-1 elements (L1M4 and L1MB3), which were either nonexistent or barely visible in control samples (**Fig. 5A and 5B**). These junctions facilitate the insertion of a novel exon between canonical exons (Exon 1 and Exon 2) that is derived from LINE-1 sequences. The incorporation of this cryptic exon into the mature transcript, leading to the generation of alternative isoforms, is further confirmed by junction level evidence (J1, J2, J3) (**Fig. 5C**).

**Figure 4.**
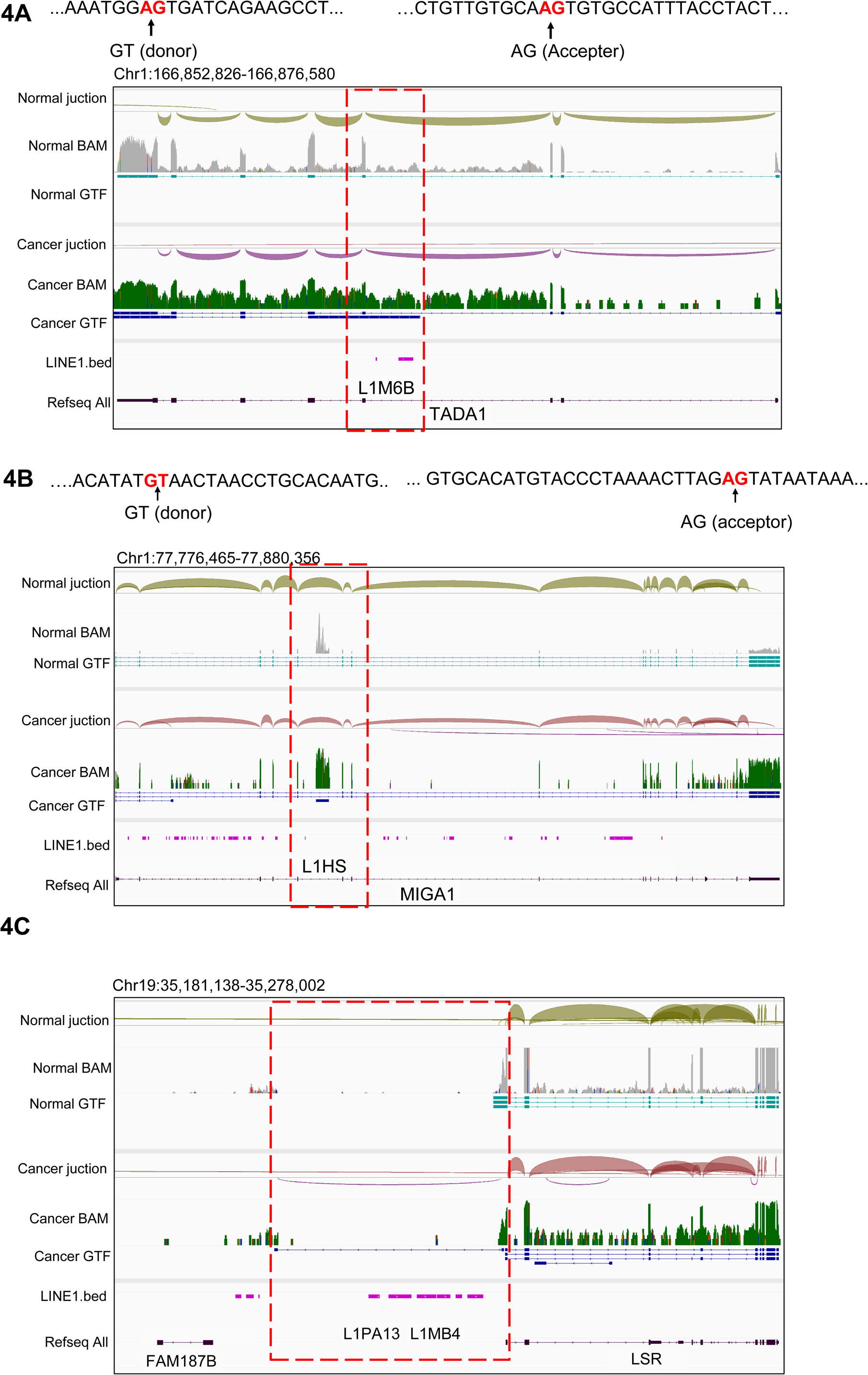
Genome browser visualization of LINE-1 associated exonization events in NSCLC. (A) Integrative genome browser view of a representative locus on chromosome 1 (Chr1:166,852,826-166,876,580) comparing normal and tumor samples. Tracks show splice junctions, RNA-seq coverage (BAM), transcript annotations (GTF), LINE-1 elements (LINE1.bed), and RefSeq gene models. Tumor samples display increased read coverage and the appearance of novel splice junctions within an intronic region containing a LINE-1 element. The highlighted region (red dashed box) marks a cryptic splice site overlapping LINE-1, consistent with LINE-1 mediated exonization. (B) Genome browser view of a second locus on chromosome 17 (Chr17:77,776,465-77,880,356). Compared to normal samples, tumor samples exhibit enhanced RNA-seq signal and multiple novel splice junctions. These junctions map to regions enriched for LINE-1 insertions, as indicated by the LINE1 track. The highlighted region shows LINE-1 associated splicing events. (C) Genome browser view of a third locus on chromosome 19 (Chr19:35,181,138-35,278,002). Tumor samples show strong read enrichment and aberrant splice junction formation within a LINE-1 rich intronic region, whereas normal samples show minimal or no such events. The highlighted region indicates inclusion of LINE-1 derived sequence into the transcript.

**Figure 5.**
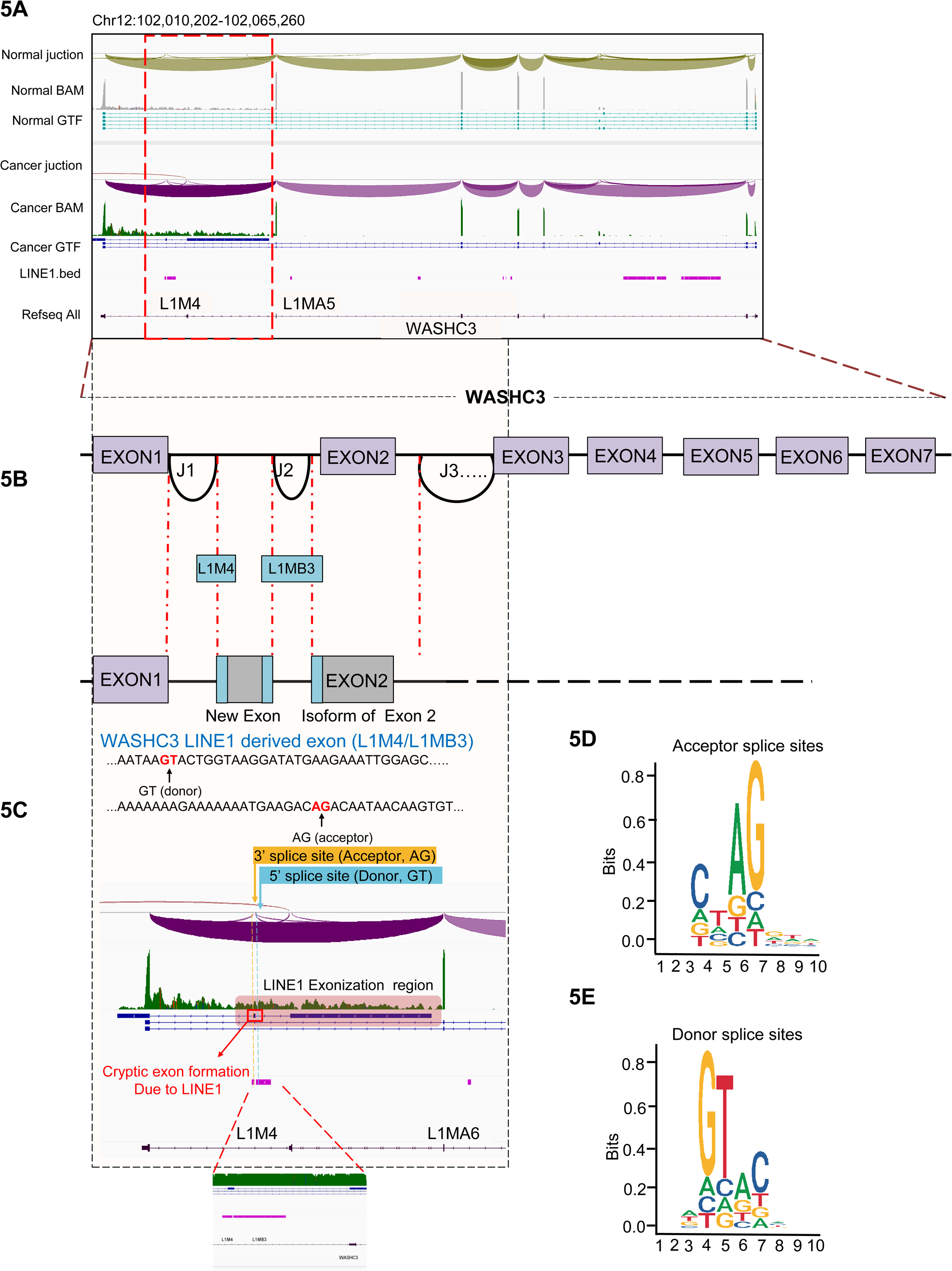
LINE-1 driven exonization generates cryptic exons in the WASHC3 locus. (A) Integrative genome browser view of the WASHC3 locus (Chr12:102,010,202-102,065,260) comparing normal and tumor samples. Tracks display splice junctions, RNA-seq coverage (BAM), transcript annotations (GTF), LINE-1 elements (LINE1.bed), and RefSeq gene models. Tumor samples show enhanced read coverage and novel splice junctions within an intronic region containing LINE-1 elements, whereas normal samples lack these features. The highlighted region (red dashed box) marks a LINE-1 associated cryptic splicing event. (B) Schematic representation of LINE-1 mediated exonization in WASHC3. Intronic LINE-1 elements (L1M4 and L1MB3) introduce cryptic splice donor and acceptor sites, leading to the formation of a novel exon between canonical exons. Multiple splice junctions (J1, J2, J3) contribute to alternative isoform generation. (C) Sequence level view of the exonization event showing precise splice junctions within the LINE-1 sequence. Canonical splice signals are identified, including 5′ donor (GT) and 3′ acceptor (AG) motifs. The highlighted region indicates the LINE-1 derived exon, confirming that cryptic splice sites embedded within LINE-1 drive exon inclusion in tumor samples. (D) Sequence logo of acceptor splice sites (3′ splice sites) derived from LINE-1 exonization events. The motif shows strong enrichment of the canonical AG dinucleotide, along with conserved upstream nucleotide preferences. (E) Sequence logo of donor splice sites (5′ splice sites) associated with LINE-1 exonization. The motif displays the conserved GT dinucleotide, consistent with canonical splice donor signals.

The LINE-1 sequence sequences provide the splice donor (GT) and splice acceptor (AG) site, thus enabling the splice machinery to identify the exonization site, based on an annotation of the exonization region. The validity of the exonizations was demonstrated by analyzing the sequence features of the splice junctions, where a highly enriched representation of canonical splice site signals was observed, consisting of conserved AG acceptor sites and GT donor sites (**Fig. 5D and E**). Notably, in the tumors, the visualization of IGV images demonstrated increased coverage and splice junction signals at the LINE-1 site, providing strong evidence for genuine exonization events. These exonizations promote isoform diversity of the WASHC3 gene by modifying its transcript structure through novel exon formation and exon elongation. To understand the nucleotide composition of the LINE-1-derived region in the WASHC3 gene region (chr12:102017647-102018468), we analyzed the LINE-1 sequence composition. Consistent with previous observations, canonical splice sites were identified in the LINE-1 element (L1M4/L1MB3). Importantly, the sequence had a canonical 5’ splice donor site (GT). These observations indicate that the exonized region harbors active splice donor and acceptor sites, facilitating recognition and utilization by the splice machinery. Comparable features suitable for splicing were also identified at other loci, which reinforces this conclusion. For example, the presence of short splicing motifs (“TT”) in the LINE-1 associated region was observed at the MIGA1 locus, while TADA1 exhibited normal splicing motifs (donor: “AG”, acceptor: “GA”). These results provide additional evidence that cryptic splicing sites exist in the LINE-1 element. Overall, these results indicate that LINE-1 elements present in intronic regions serve as splice competent sites, resulting in the generation of new exons and alternatively spliced transcripts in cancer. In summary, these results support a more general hypothesis where cryptic splicing sites embedded in transposons are utilized by exonization through LINE-1s to remodel gene transcripts.

## Discussion

We present a t transcriptome wide and locus-level analysis of LINE-1 mediated exonization in NSCLC. Demonstrating its importance as a mechanism of transcriptome remodeling in cancer. Our results show that LINE-1 elements function as a functional reservoir of splice competent sequences, contributing to isoform diversity, by combining expression profiling, splice junction analysis, recurrence mapping, and sequence level validation. We found a general pattern of transcriptional upregulation in NSCLC, including increased expression of important oncogenic drivers like EGFR, KRAS, MET, and PIK3CA, which is consistent with LINE-1 reactivation in cancer (Chevallier et al., 2021). Additionally, correlation analysis showed a significant correlation between LINE-1 expression and a number of classical oncogenes, indicating a connection between oncogenic transcriptional programs and LINE-1 activity. However, locus level IGV analysis did not show direct LINE-1 mediated exonization within these conventional oncogenes despite this association, suggesting that indirect or upstream mechanisms rather than direct exon inclusion may be responsible for their regulation. On the other hand, we found that LINE-1 elements are directly implicated in exonization events in a number of non-canonical and poorly described genes (such as WASHC3, MIGA1, LSR and TADA1). These genes play functions in vesicular trafficking, mitochondrial dynamics, and transcriptional control, which makes them functionally important to cancer even if they are not traditional oncogenic drivers. This implies that rather than being limited to well-known oncogenes, LINE-1 mediated exonization preferentially targets a larger network of genes that either directly or indirectly contribute to tumor biology. The ubiquitous occurrence of LINE-1 mediated exonization in NSCLC samples, with repeated occurrences seen at particular loci, is a significant finding of our investigation. Exonization happens in both sense and antisense orientations, with a preponderance of antisense derived events, according to strand-specific studies. This is in line with the enrichment of splice compatible motifs in antisense LINE-1 sequences. Our findings show that LINE-1 elements contain canonical splice donor (GT) and acceptor (AG) sites, which the splicing machinery can use to create new exons. Both sequence-level analysis and visualization by IGV have confirmed the validity of this. As can be seen at loci such as WASHC3, the process of exonization causes the formation of new exons and transcript variability. It is important to note that our findings validate a theory wherein regulation by LINE-1 is heavily dependent on the genomic context and locus. The functionality of the effect will depend on where the LINE-1 sequence lies within the genome, its orientation, and local splicing environment. The selective integration of the LINE-1 sequences into certain genes but not others, including classical oncogenes, may be due to this dependence.

In conclusion, our study shows that exonization by LINE-1 is a prevalent and functionally relevant pathway for transcriptome diversity in NSCLC. Exonization by LINE-1 sequences increases the diversity of transcripts without changing any genomic sequence by acting as a reservoir of splice sites. These findings illustrate LINE-1 exonization as a potential determinant of tumor heterogeneity and evolution and provide novel insights into the role of transposons in altering gene structure. To determine the functional ramifications of certain exonization events, such as their influence on cellular phenotypes, protein function, and clinical outcomes, further research will be needed. Further research is necessary to determine whether LINE-1 exonization events can be used as biomarkers or therapeutic targets.

## Material and methods

### Bulk RNA-seq processing

Bulk RNA-seq data were first aligned to the human reference genome (GRCh38) using STAR (Dobin et al., 2013) in two-pass mode to improve splice junction detection in order to identify LINE-1 exonization events. In order to prepare the alignments for further analysis, they were sorted into BAM files. RegTools (Feng et al., 2018) was then used to extract splice junctions from the BAM files, and BEDTools (Quinlan, 2014) was used to intersect junction BED files with LINE-1 genomic annotations in order to find junctions that overlapped LINE-1 elements. Candidate junctions were screened based on a minimum read support threshold of five reads, overlap with annotated LINE-1 families (such as L1HS, L1PA2, L1PA3), and elimination of duplicate junctions based on genomic coordinates in order to guarantee robustness. The GENCODE v38 GTF file was used to annotate filtered junctions. Genomic ranges were used to calculate overlaps between LINE-1 locations and gene characteristics (exon, CDS, UTR). Exonization was characterized as splice junctions connecting LINE-1 sequences to host gene transcripts. Events were categorized into chromosomal categories (exonic, intronic, UTR, and intergenic). Gene expression matrices were constructed from bulk RNA-seq count data. Counts were normalized using log2 transformation (log2(count + 1)) for downstream visualization and correlation analysis. Key oncogenic genes relevant to NSCLC, including EGFR, KRAS, NRAS, BRAF, PIK3CA, MET, and CTNNB1,WASCH3,TADA1,MIGA1 were selected for focused analysis.

### Quantification of LINE-1 exonization events

The number of distinct splice junctions that spanned annotated LINE-1 elements was used to quantify LINE-1 exonization events for each sample. Only high-confidence junctions those with enough read support and non-duplicated genomic coordinates were kept in order to guarantee accuracy. After that, event counts were combined at several levels: at the subfamily level to differentiate between several LINE-1 families, such as L1HS and L1PA, and at the sample level to allow comparisons across experimental circumstances. We were able to evaluate the overall and lineage-specific contributions of LINE-1 elements to transcriptome diversity thanks to this stratification. Low-confidence junctions were eliminated in order to minimize background noise and guarantee that only strong events were taken into account. In order to assess variations in LINE-1 activity across biological states, the resultant counts were compared across normal and tumor samples after being utilized to calculate exonization load, which provided a quantifiable measure of LINE-1 derived splicing activity.

### Correlation analysis between LINE-1 and gene expression

The number of distinct splice junctions that spanned annotated LINE-1 elements was used to quantify LINE-1 exonization events for each sample. Only high-confidence junctions those with enough read support and non-duplicated genomic coordinates were kept in order to guarantee accuracy. After that, event counts were combined at several levels: at the subfamily level to differentiate between several LINE-1 families, such as L1HS and L1PA, and at the sample level to allow comparisons across experimental circumstances. We were able to evaluate the overall and lineage specific contributions of LINE-1 elements to transcriptome diversity thanks to this stratification. Low-confidence junctions were eliminated in order to minimize background noise and guarantee that only strong events were taken into account. The resulting counts were used to compute exonization burden, providing a quantitative measure of LINE-1 derived splice activity, and were subsequently compared between normal and tumor samples to evaluate differences in LINE-1 activity across biological states. In order to assess variations in LINE-1 activity across biological states, the resultant counts were compared across normal and tumor samples after being utilized to calculate exonization burden, which provided a quantifiable measure of LINE-1 generated splicing activity.

### L1 and Refseq reference

The repeat gene family annotation for hg38 was downloaded from the RepeatMasker database and repeat families annotated as “L1” were extracted for further analysis. We then selected the L1 families whose coordinates located within the upstream and downstream 50 kb of coding genes from NCBI Refseq database. The sequences of these L1 families were extracted from the human genome hg38 using bedtools getfasta based on their genome coordinates. Make blastdb of blast 2.2.26 + was used to build the reference for blast mapping. Meanwhile, all the repeat sequences from RepeatMasker were also extracted and genome Generate was used to build the reference for STAR mapping to remove false positive chimeric candidates.

Refseq RNA sequences was downloaded from NCBI data base. Repeatmasker 4.0.7 (http://www.repeatmasker.org >) was used to identify and mask the repeat sequences in the RNA sequence using hmmer method. The repeat sequence identified from the RNA sequence was replaced with Ns. Two reference transcriptomes were generated with makeblastdb of blast 2.2.26 +, one with original Refseq sequence and one for masked reference. A bwa reference for the unmasked RNA sequence was also built with bwa index with default parameters.

### Identification of differentially expressed L1 chimeric transcripts Events

Raw L1 chimeric transcripts sample counts were normalized with the total RNAseq reads in each sample and then log transferred. Then two strategies were used to identify differentially expressed L1 chimeric transcripts events. 1) limma test was performed on the normalized expression value of L1 chimeric transcripts in tumor samples against normal controls. The gene chimeric transcripts events with L1 chimeric *p* value < = 0.05, log2 fold change > 0 were selected as differentially expressed events. 2) For each event, the number of occurrences in cancer and control of gene chimeric transcripts and normal transcripts were counted and Fisher’s exact test was performed to calculate the significance p value. The gene chimeric transcripts events with *p* value < = 0.05 were selected as differentially expressed events. Finally, the union set of differentially expressed events identified in the two strategies were selected as the differentially expressed events.

## Resource Availability

The RNA-seq and datasets used in this study were obtained from the public repository NCBI Gene Expression Omnibus (GEO) under the accession number GSE40419.

## Author contribution

ASP: methodology, investigation, formal analysis; ASP and AK: Writing original draft; BT: conceptualization, fund acquisition, and final editing.

## Declaration of Interests

The authors declare no competing interests.

## Funding

This work was supported by the DBT/Wellcome Trust India Alliance fellowship (IA/I/22/2/506501) to Bhavana Tiwari (BT). ASP received SRF support from IA Fellowship to BT.

## Notes

### Competing Interest Statement

The authors have declared no competing interest.

